# Proactive Coral Reef Restoration Using Thermally Tolerant Corals in Hawai‘i

**DOI:** 10.1101/2025.03.28.646055

**Authors:** Hanalei Ho‘opai-Sylva, Carlo Caruso, Spencer Miller, Joshua R. Hancock, Matthew Parry, Kira Hughes, Crawford Drury

**Author notes:** **Correspondence:** Crawford Drury, 46-007 Lilipuna Rd, Kāne‘ohe HI, 96744. **Author Contacts:** Hanalei Ho‘opai-Sylva Carlo Caruso Spencer Miller Joshua R. Hancock Matthew Parry Kira Hughes Crawford drury. **Data Availability Statement** All data and code needed to reproduce this analysis is available at github.com/CarloReef/RWR_Airport.

## Abstract

Effective conservation of degraded ecosystems requires mitigation of the original cause of decline, but this step can be difficult in the context of global climate change. On coral reefs, persistent environmental stress which causes coral bleaching may be addressed by using coral restoration stock which is naturally more resilient, often termed “proactive restoration” in terrestrial management. To explore the feasibility and consequences of this approach, we outplanted 391 colonies of 7 species of reef-building coral designated as ‘thermally tolerant’ or ‘thermally sensitive’ during stress testing and monitored them for 2 years using photogrammetry to evaluate tradeoffs and return-on-effort. We found no growth, complexity or effort tradeoffs when using thermally tolerant corals, but tolerant corals had lower survivorship during our monitoring period, driven primarily by one genus. These data illustrate nuanced tradeoffs and consequences to proactive reef restoration and suggest that the potential benefits of this approach may only be fully realized during future coral bleaching events.

## Introduction

We acknowledge the waters of Māmala Bay as an important place for Native Hawaiians of the past, present, and future. We hope to honor the cultural importance of this space, and the reef within, in our research.

Coral reefs create the structural complexity that supports the world’s most diverse marine ecosystems which protect shorelines from coastal erosion (Reguero et al., 2018). Coral reefs in Hawai’i are estimated to span nearly 410,000 acres across the archipelago and are ecologically and socially important (Rodgers & Cox, 2003). In addition to being an important food and income source for local families (Friedlander et al., 2005), the ko’a (coral) is considered the origin of life and is regarded as an ancient ancestor of the Hawaiian people (Gregg et al., 2015). Hawaiians trace their genealogical lineage to ko’a in the Kumulipo, a cosmogonic genealogy of the Hawaiian people, establishing a deep connection to the natural world and guiding stewardship of marine ecosystems (Kame’eleihiwa, 1992).

Native Hawaiians have utilized coral reefs to harvest food for family and friends across generations; a deeply-rooted cultural practice connecting Hawaiians to their land. The tradition of deep observation of natural processes is valued in Hawaiian culture as a means to understand resources, forecast future conditions, and plan actions, especially in coastal ecosystems. Despite the importance of coral reefs in Hawai’i, there are few local examples of active restoration, potentially due to local conditions like strong wave action, steep bathymetry, biogeographic isolation, igneous substrate and slower growing coral species (Forsman et al., 2018; Friedlander et al., 2005).

Coral reefs face a complex landscape of anthropogenic stress (Hughes et al., 2017). Among these stressors, increasing ocean temperatures are the most certain to persist and intensify, leading to coral bleaching (Brown, 1997), compromised growth (Figueiredo et al., 2022), and reproduction (Fisch et al., 2019), and eventual mortality (Baird & Marshall, 2002). Bleaching events are forecasted to become more frequent and severe (van Hooidonk et al., 2016), likely contributing to reef degradation across tropical oceans in the next several decades (Hoeke et al., 2011). Despite the threats posed by anthropogenic climate change, variation in bleaching and thermal tolerance is ubiquitous (Drury, 2020); certain populations, species and individuals are predisposed to survive under future ocean conditions (Drury, Martin, et al., 2022; Logan et al., 2014; Palumbi et al., 2014).

In an effort to combat the decline of reef habitats, coral restoration has grown in popularity as an active attempt to return reefs to a functional state using corals from a separate site (Boström-Einarsson et al., 2020; Rinkevich, 2014), typically propagated in an *in situ* or land-based nursery (Rinkevich, 1995). Reef restoration aims to assist and accelerate the recovery of reefs, protect endangered coral species, and reestablish reefs to functional and self-sustainable states (Hein et al., 2017), but if the coral stock used for restoration is not viable under current or future conditions, the conservation impacts will likely be negligible.

Despite the popularity of reef restoration, there is little systematic evidence for the long-term success of this approach, in part because projects infrequently address the underlying causes of initial decline or conduct appropriate long-term monitoring. In the context of global climate change, addressing these shortcomings is a major obstacle viewed as nearly impossible on a local scale, but is a necessary component of long-term restoration success (Shaver et al., 2022; van Oppen et al., 2017) when ocean temperatures are virtually certain to increase, even if emissions stopped today (Mauritsen & Pincus, 2017). One option to address this mismatch between environmental conditions and coral traits on reefs degraded by coral bleaching is to use restoration stock that is naturally more resilient to climate change (Caruso et al., 2021) with the expectation that such corals contribute to ‘pre-adaptation’ or genetic rescue of coral populations on the reef(Bay et al., 2017; Shaver et al., 2022). While ecological restoration is inherently a “reactive” activity (Hein et al., 2021), approaches that focus efforts on preparation for future conditions may be termed “proactive restoration” or “preemptive restoration” (Muzika, 2017; Schweiger et al., 2019; Schweitzer et al., 2014). Experimental designs that outplant corals that can withstand warm and increasing ocean temperatures while providing durable coral cover, species diversity, and structural complexity may lead to more efficient conservation efforts and should be considered in future restoration programs (Caruso et al., 2021; Morikawa & Palumbi, 2019). However, to responsibly implement these changes requires a thorough understanding of tradeoffs between thermal tolerance and other important traits under stressful and benign conditions, including growth, structural complexity, survivorship, fecundity, corallivore palatability, and many others. If the most thermally tolerant corals in a population are selected for restoration but do not contribute effectively to the structure and function of a coral reef, this solution to a biological-environmental mismatch may be ineffective, which is why these techniques must be thoroughly explored (Caruso et al., 2021).

A second major obstacle to successful reef restoration is establishing a monitoring program which collects “meaningful, consistent, comparable, and quantitative data” to inform decision makers and compare outcomes across projects (*Coral Reef Restoration Monitoring Guide: Methods to Evaluate Restoration Success from Local to Ecosystem Scales*, n.d.; Hein et al., 2017; Platz et al., 2022). Short timelines (< 1 year), imprecise techniques (visual surveys, transects, photo quadrats, chain-and-tape), and limited financial or logistical support are commonly identified challenges for monitoring coral restoration projects (Bellwood et al., 2019; Boström-Einarsson et al., 2020). One approach to improve monitoring throughput, reproducibility and quality is the use of Structure-from-Motion (SfM) photogrammetry, a scalable, non-invasive method for monitoring a suite of outcomes (e.g., bleaching (Yadav et al., 2023), settlement (Barrows et al., 2023), growth (Lange et al., 2022) and structural complexity, (Miller et al., 2021) at restoration sites. SfM photogrammetry removes the time constraint on many of the most important data collection steps (e.g., measurements, species identification), which can be completed after the fact, increasing the quality and quantity of data collected from the field (Burns et al., 2015; Ferrari et al., 2021; Fukunaga et al., 2022).

Overcoming these obstacles to collect accurate data over ecologically-relevant timescales remains a challenge, but is critically important because it facilitates the assessment of effectiveness within and between restoration projects and programs using tools such as Relative Return-on-Effort (Suggett et al., 2019). RRE calculations allow for the explicit comparison of cost, practicality and scalability of restoration efforts by evaluating growth and survivorship of outplanted corals from restoration programs around the world (Henry et al., 2023; Howlett et al., 2021). This framework allows practitioners to develop metrics of success and offers an objective approach to evaluate performance of new and ongoing projects. With the growing collection of quantitative data on restoration practices available today, the ability to understand which approaches deliver the best outcomes can inform best practices for site-specific locations where little to no local history of restoration exists (Boström-Einarsson et al., 2020).

Here we integrate the concepts of proactive reef restoration with long-term monitoring and effort measurement in a research-scale restoration project in Hawai’i. We used short-term stress testing on experimental biopsies to assay thermal tolerance before establishing restoration plots of thermally tolerant and thermally sensitive corals of opportunity on the south shore of O’ahu. We monitored these outplants for attachment, growth, survivorship, and reef complexity using SfM to assess the consequences and effectiveness of selecting thermally tolerant coral stocks for restoration and compared the results to global restoration outcomes using RRE calculations to evaluate the potential for reef restoration in Hawai’i.

## Methods

Corals of opportunity collected in response to a 2018 ship grounding on the south shore of O’ahu were transplanted in a nearby *in situ* nursery to be used as restoration stock. In 2020, we collected replicate biopsies from stock corals of 9 species and used a 2-week heat stress capable of resolving natural bleaching phenotypes (Caruso et al., 2025; Drury, Dilworth, et al., 2022) to designate corals as thermally tolerant or thermally sensitive based on within-species performance in the heat assay. We designated 6-meter diameter plots at a nearby forereef between 8.2m to 14.5m depth and randomly designated plots as ‘high tolerance’ (n=8), ‘low tolerance’ (n=9), or ‘control’ (n=6; no outplanting). A total of 391 corals (n=23 to 25 colonies per plot) were randomly assigned to non-control plots based on thermal tolerance designation, creating replicate plots with exclusively thermally tolerant corals, exclusively thermally sensitive corals, or no outplanted corals (control).

All sites were monitored with Structure from Motion (SfM) photogrammetry before outplanting and periodically over 2 years; we derived attachment rates, survivorship, growth (surface area) and structural complexity from SfM data. These data were used to calculate Relative Return on Effort (RRE) scores for comparison with other restoration programs (Suggett et al., 2019). Detailed methods including outplanting design, stress-testing, data collection and processing, and statistical analysis are described in the Supporting Information.

## Results

### Coral Plots

Our selected plots had an average of 79.2 ± 16.3% (mean± 1SD) hard coral cover before any outplanting occurred. We outplanted 391 coral colonies across 17 plots. Of these, 334 colonies (112 *Montipora capitata*, 12 *Montipora patula*, 14 *Pocillopora meandrina*, 18 *Porites compressa*, and 186 *Porites lobata*) had usable data after model reconstruction and data quality control. Outplanted corals including *Porites evermanni, Montipora flabellata, Pocillopora grandis*, and *Pavona varians* were excluded from downstream analysis because there were fewer than 10 total colonies outplanted for each species. Between 23 and 26 corals were outplanted at each of the 17 plots (9 thermally sensitive plots and 8 thermally tolerant plots; Figure 1C). Across all plot photogrammetry models there was an average of 1.7 pixel root-mean-square (RMS) reprojection error, 0.39 mm linear error (0.66%), 0.11 mm^2^ area error (1.5%), and 0.02 mm^3^ volumetric error (2.5%). We also monitored a total of 150 corals that were randomly selected in 6 control plots where no corals were outplanted (25 corals x 6 plots), 146 remained after model reconstruction and data QC.

**Figure 1.**
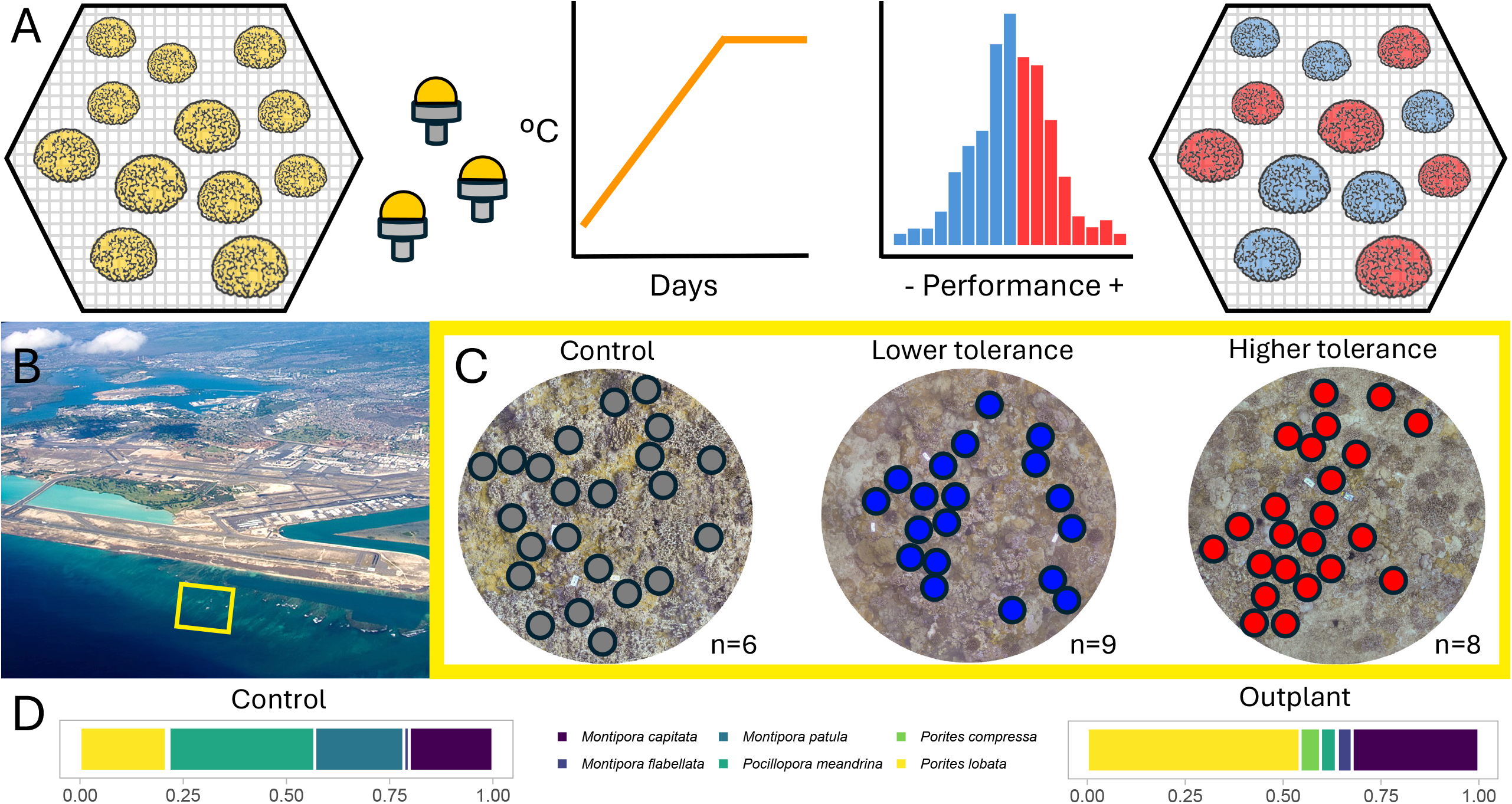
Schematic representation of coral collection and experimental design. (A) Biopsies were taken from Corals of Opportunity (COO) on NOAA’s *in-situ* nursery table located south of the Honolulu Airport and stress tested using a 2 week thermal exposure and performance was used to assign thermal tolerance to original colonies. (B) The restoration site was divided into 23 study plots adjacent to the nursery on South Shore, O’ahu (photo reproduced with permission from Gabor Hajdufi). (C) COO (n=23-25 per plot) were outplanted according to thermal tolerance assignment. For control plots, corals were picked from orthomosaic imagery *post hoc*. N values represent number of plots, point color represents plot type. (D) Proportion of species at control plots compared to outplant plots.

### Plot Complexity

Outplanting took place 3 months after initial complexity data of plots was collected. Outplanting corals significantly increased the fractal dimension of plots (p=0.002; Figure 2A) and all but one plot had an increased fractal dimension after outplanting. Fractal dimension continued to increase over time in both plot types (p=0.009; Figure 2B), although the interaction between plot type and time was not significant (p=0.076). Examining outplant plots only (i.e., excluding control plots), fractal dimension significantly increased over time (p=0.003, Figure 2C), but there was no significant interaction between time and thermal tolerance level on complexity (p=0.906, Figure 2C).

**Figure 2.**
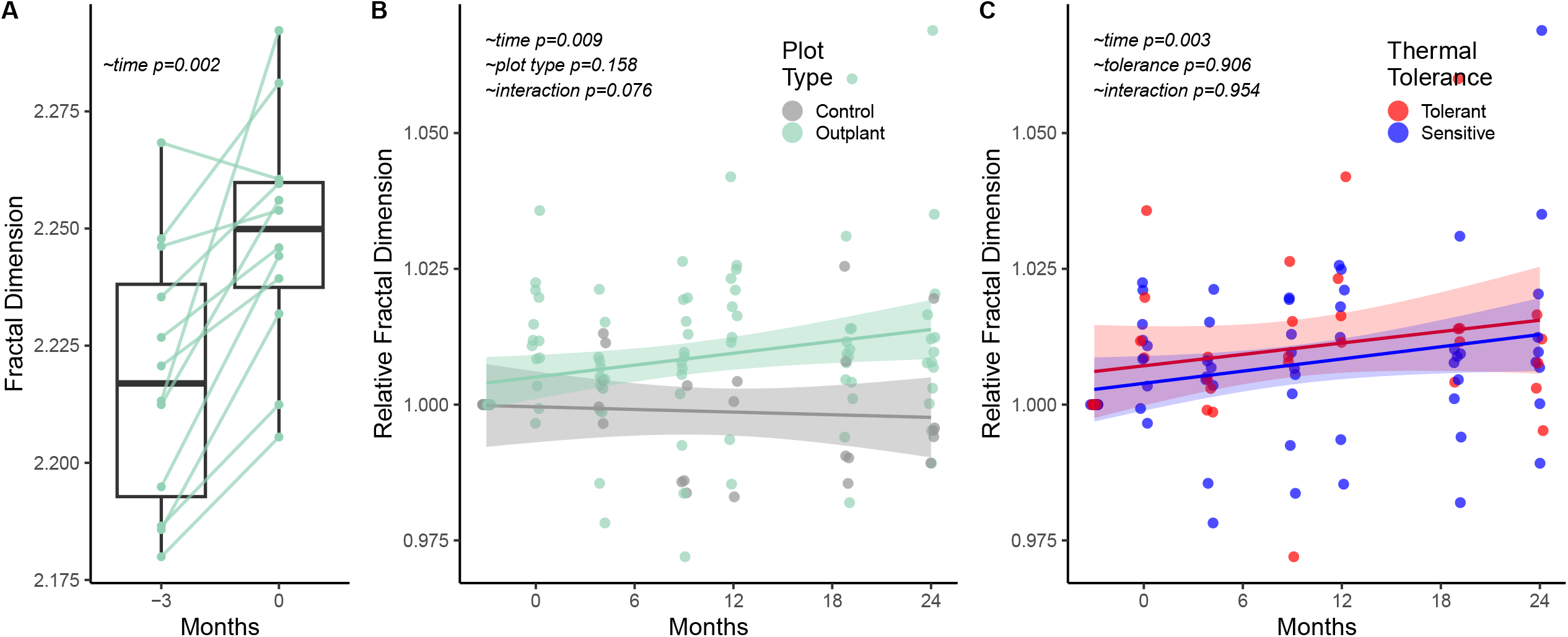
Measure of structure and complexity of restoration plots. Initial plot complexity was measured 3 months before outplanting. (A) Raw fractal dimension of the reef measured before (timepoint 1, -3 months) and after (timepoint 2, 0 months) coral outplants were added to a plot. Each point represents a plot, connected between timepoints. Box represents 25-75 quartiles, whiskers represent 1.5IQR. (B) Relative fractal dimension of outplanted and control plots over time, where color represents plot type. Raw fractal dimension data was used for statistical analysis. Shading is 95%CI. (C) Relative fractal dimension of thermally tolerant and sensitive plots over time, where color represents plot type. Shading is 95%CI. Raw fractal dimension data was used for statistical analysis.

### Attachment, Survivorship, and Growth

Attachment probability of outplanted corals was significantly lower than control corals (p<0.001; Figure 3A); after 2 years, 97% of control corals and 74% of outplanted corals remained attached (Figure 3B). Among corals that remained attached, survivorship probability of outplanted corals was significantly lower than control corals (p<0.001; Figure 3C); at the 2 year mark, 97% of attached control corals and 82% of attached outplanted corals remained alive (Figure 3D).

**Figure 3.**
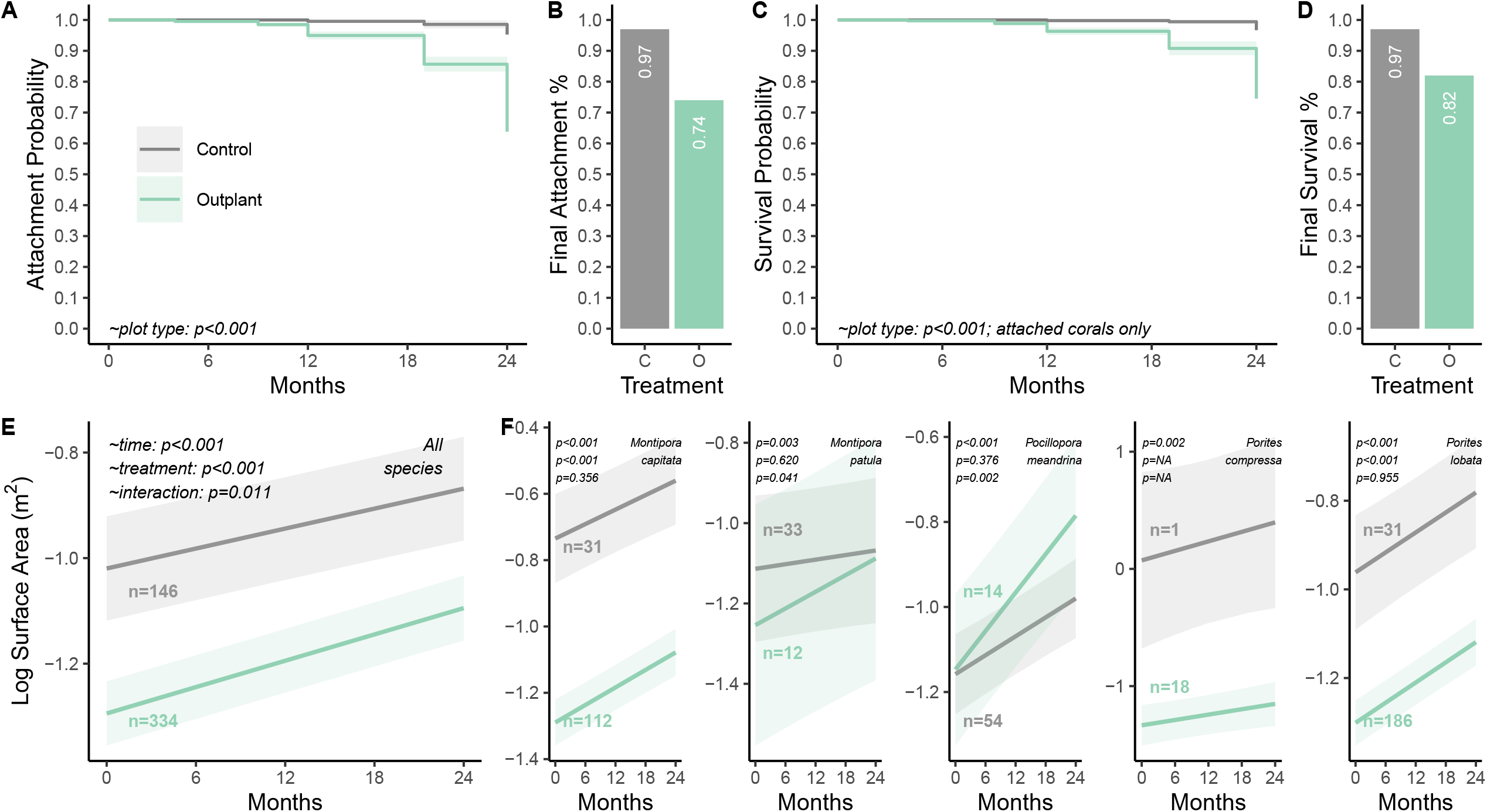
Attachment, survivorship and growth of outplanted and control corals. (A) Attachment probability of control and outplanted corals over 24 months. (B) Final attachment percentage of control and outplanted corals. (C) Survival probability of corals that remained attached. (D) Final survival percentage of corals that remained attached. (E) Growth of outplant and control coral colonies over time, measured in m^2^. (F) Species-specific growth. Colors represent control and outplanted corals. Surface area plots are visualized using the model output. Shading represents 95%CI throughout the figure.

Across all species, control corals were significantly larger than outplanted corals (p<0.001) throughout the observation period, the size (m^2^) of both control corals (n=146) and outplanted corals (n=334) significantly increased over time (p<0.001), and there was a significant interaction between time and treatment (p=0.011), with outplanted corals growing at a faster rate (Figure 3E).

*Montipora capitata* significantly increased in size over time (p<0.001), control corals were significantly larger than outplanted corals (p<0.001) and there was no significant interaction between time and treatment on coral size (p=0.356; Figure 3F). *Montipora patula* significantly increased in size over time (p=0.003), control corals were not a significantly different size than outplanted corals (p=0.620) and there was a significant interaction between time and treatment on coral size (p=0.041), where outplanted corals grew faster (Figure 3F). *Pocillopora meandrina* significantly increased in size over time (p<0.001), control corals were not a significantly different size than outplanted corals (p=0.376) and there was a significant interaction between time and treatment on coral size (p=0.002), where outplanted corals grew faster (Figure 3F). *Porites compressa* was poorly represented in the control (n=1), so we excluded it from part of the analysis, but corals significantly increased in size over time (p=0.002; Figure 3F). *Porites lobata* significantly increased in size over time (p=0.003), control corals were significantly larger than outplanted corals (p<0.001) and there was no significant interaction between time and treatment on coral size (p=0.955; Figure 3F).

### Thermal Tolerance Impacts on Survivorship and Growth

Thermally tolerant corals had a 189% higher hazard, translating to significantly lower survival probability than thermally sensitive corals (p<0.001; Figure 4A). The addition of species (p<0.001) to the model significantly impacted survivorship (Figure 4A). At the end of 24 months, 77% of thermally tolerant corals and 87% of thermally sensitive corals survived (Figure 4B).

**Figure 4.**
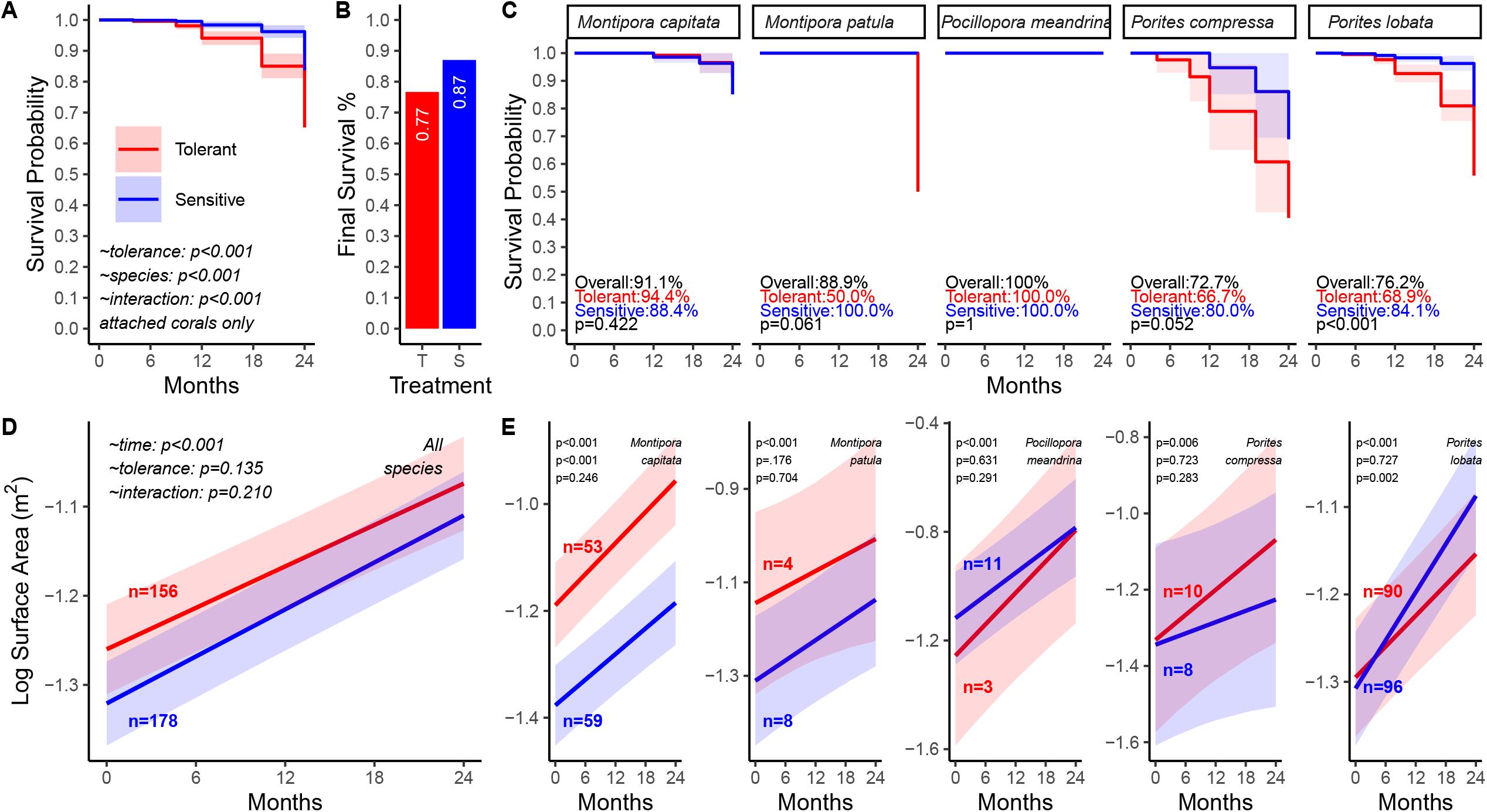
Attachment, survivorship and growth of thermally tolerant and sensitive corals. (A) Survival probability of thermally tolerant and thermally sensitive corals. (B) Final survival percentage of thermally tolerant (T) and sensitive (S) corals. (C) Survival probability of thermally tolerant and sensitive corals separated by species. (D) Growth of tolerant and sensitive outplants over time measured by surface area (m^2^). (F) Species-specific growth. Colors represent tolerant and sensitive corals. Surface area plots are visualized using the model output. Shading represents 95%CI throughout the figure.

Within species, thermally tolerant *Porites lobata* had significantly lower survivorship (p<0.001), while thermally tolerant *Montipora patula* and *Porites compressa* had lower survival rates that were marginally insignificant (p=0.061 and p=0.052; Figure 4C). *Montipora capitata* and *Pocillopora meandrina* had nearly identical survival rates between tolerance groups (Figure 4C).

Across all species, the size (m^2^) of both tolerant corals (n=156) and sensitive corals (n=178) significantly increased over time (p<0.001), there was no significant difference in size (p=0.135) and there was no significant interaction between time and tolerance (p=0.210; Figure 4D).

In *M. capitata*, corals significantly increased in size over time (p<0.001), and tolerant corals were significantly larger than sensitive corals (p<0.001), but there was no significant interaction between time and thermal tolerance on size (p=0.246; Figure 4E). *Montipora patula* significantly increased in size over time (p<0.001), but there was no difference in size between tolerant and sensitive corals (p=0.176) and no significant interaction (p=0.704; Figure 4E). *Pocillopora meandrina* significantly increased in size over time (p<0.001), but there was no difference in size between tolerant and sensitive corals (p=0.631) and no significant interaction (p=0.291; Figure 4E). *Porites compressa* significantly increased in size over time (p=0.006), but there was no difference in size between tolerant and sensitive corals (p=0.723) and no significant interaction (p=0.283; Figure 4E). *Porites lobata* significantly increased in size over time (p<0.001), there was no difference in size between tolerant and sensitive corals (p=0.727), but there was a significant interaction between thermal tolerance and time, (p=0.002; Figure 4E), where thermally tolerant corals grew more slowly.

### Evaluating Relative Return on Effort (RRE)

Growth and survivorship of outplanted corals were compared using Relative Return-on-Effort (Suggett et al., 2019), which was developed as a tool for restoration practitioners to determine which coral species and methods allow for the most successful return on restoration efforts. Although thermally sensitive corals had 8% higher RRE scores than thermally tolerant corals, this difference was not significant (p=0.519; Figure 5AB). While RRE scores were significantly different across 7 global regions (p<0.001), the average RRE score in Hawai’i was 10.8, which was not significantly different from the global average of 11.5 (p=0.351; Figure 5CD). Outplanted corals in Hawai’i generally had similar survivorship and lower growth than restoration projects in other regions (Figure 5D).

**Figure 5.**
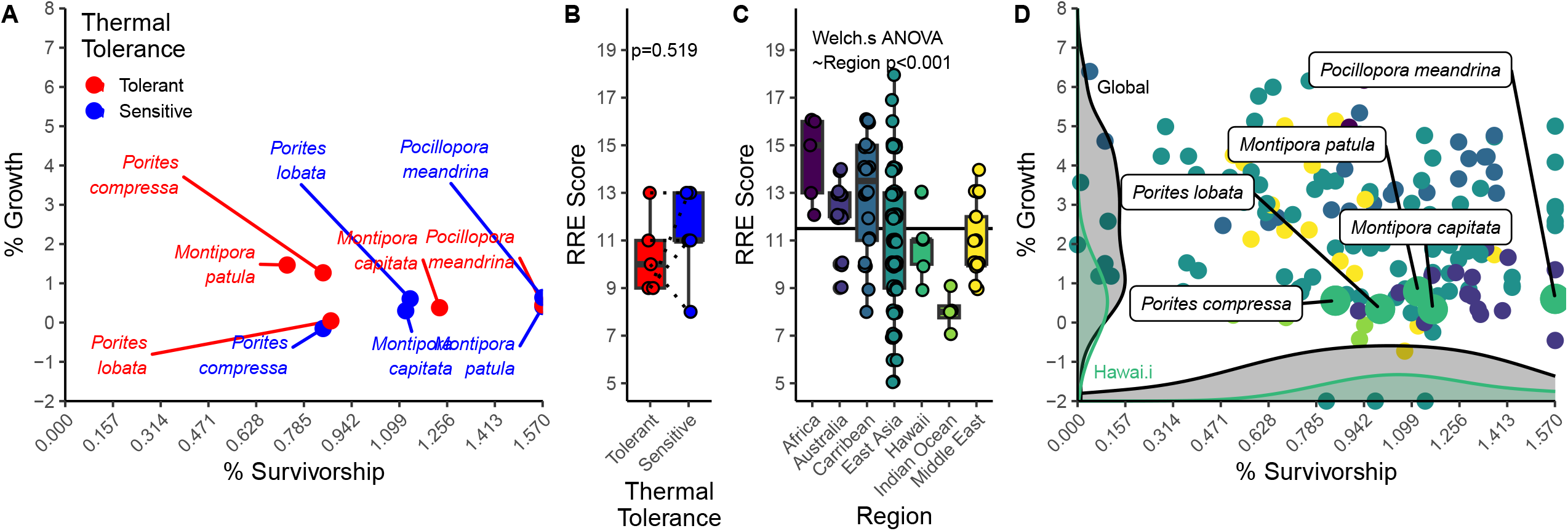
Relative Return on Effort (RRE) of coral outplants. (A) Growth and survivorship of the five primary species analyzed in this project. Colors represent thermal tolerant assignments. (B) Boxplot of RRE scores of thermally tolerant and sensitive corals in Hawai’i (this study). (C) Boxplot of RRE scores of global restoration projects. (D) Transformed growth and survivorship of global restoration projects compiled from (Henry et al., 2023; Howlett et al., 2021; Suggett et al., 2019). Colors represent regions. Density plots represent growth (y-axis) and survivorship (x-axis) in Hawai’i (green, this study) and global projects (black). Box represents 25-75 quartiles, whiskers represent 1.5IQR.

## Discussion

Efforts to protect, conserve, and restore reefs are crucial, especially in Hawai’i and other regions where community wellbeing is strongly tied to reef health. Successful conservation work can perpetuate cultural practices related to natural resources and avert cultural dissonance arising from their loss (Juel Clemmensen, 2014), especially in Hawai’i where Indigenous Hawaiians protected coral reefs before the introduction of Western science (Kikiloi, 2003). It is essential to recognize and respect the local knowledge systems employed in Hawai’i and to acknowledge their efficacy alongside conventional Western scientific methodologies. Evidence points to the weaving of western and indigenous science in modern research as beneficial for both visiting scientists conducting research and the local community whose inherent right it is to oversee the protection of their resources (Alexander et al., 2021; Cooke & Arlinghaus, 2024). Though our activities in this research-scale restoration project were mainly informed by Western science, we recognize the importance of generational Hawaiian knowledge on natural resource management, anticipate that future coral restoration efforts in Hawai’i will be community driven, and strive to find better ways to integrate local knowledge systems into coral restoration.

Our experimental outplanting illustrates the efficacy of restoration in Hawai’i through durable attachment, high survivorship and growth of corals outplanted over 2 years. The relatively long monitoring duration, large footprint and high species diversity compared to typical projects (Boström-Einarsson et al., 2020) provide important context for reef restoration in the Central Pacific, which has historically been undertaken as a mitigation response for anthropogenic damage in commercially important, relatively low wave-energy environments (Maragos et al., 2006; Rodgers et al., 2017). Outplant attachment rates in this study (72% after two years) were higher than expected given typical wave energy conditions in Hawai’i that are higher than many other locations worldwide where coral restoration has been tested. Several factors may contribute to the difference between control and outplanted coral retention, including skeletal accretion onto the substrate (lower in outplanted corals), the higher frequency of branching and massive morphologies in outplanted colonies, and high incidence of encrusting corals in control plots, all of which favor control corals under periodic high wave energy that would be more likely to dislodge outplanted colonies (Tagliafico et al., 2018). Conversely, the introduction of outplanted corals to plots increased complexity more quickly than control plots over time and we detected faster growth rates in outplanted corals, highlighting the benefits of outplanting for structure and function, even on reasonably healthy reefs.

Outplanted corals had lower survivorship than corals natively in control plots, which may be due to factors beyond local adaptation to a home site. Although the endpoint survivorship differences were moderate between groups (15% lower in outplanted corals), a number of factors outside our experimental framework likely influenced this capacity: corals of opportunity experienced damage prior to collection, were subjected to sampling stress during biopsies (Okubo et al., 2007), and had to acclimate to new environmental conditions during movement to a common garden nursery and subsequently during transplantation into plots with changing light, water temperature, and wave action, all of which impact coral physiology (Putnam et al., 2017). Perhaps most importantly, control corals had significantly larger initial sizes, which improves survival probability (Furby et al., 2017; Madin et al., 2014). Although most outplants survived and grew, this result highlights the trade-offs between size, survivorship, diversity and the logistical trade-offs of outplanting larger corals. Overall, survivorship in this project was 82% over 2 years, higher than typical rates (∼65%) reported over shorter timespans (<1 year) (Boström-Einarsson et al., 2020), supporting the viability of restoration in Hawai’i.

Our results illustrate nuanced tradeoffs to thermal tolerance, however our monitoring did not capture a coral bleaching event, when the primary traits selected by the heat stress in this experiment are expected to manifest (Caruso et al., 2025). These data fit into a previously defined framework, where the benefits of thermal tolerance (Jones & Berkelmans, 2010) may only emerge under stressful conditions (Bay & Palumbi, 2017; Cunning et al., 2015; Ladd et al., 2017) because there is an ultimate energetic ceiling for growth, reproduction, resistance and recovery in any given coral colony (Lesser, 2013). Empirical tests, such as this study, are important for determining the place-based consequences of selecting coral stocks by thermal tolerance. Survivorship of thermally tolerant corals was significantly lower than their thermally sensitive counterparts, but this pattern was driven primarily by *Porites compressa* and *P. lobata*. Similarly, there was no difference in growth rate between thermally tolerant and sensitive corals, except in *P. lobata*. We hypothesize that genotype-environment interactions at very small spatial scales are the best explanation for these patterns in one species, especially because *Porites spp*. in Hawai’i harbor a consistent, low-diversity community of *Symbiodinacea* (primarily C15) relative to other species (Stat et al., 2013). This result highlights the importance of species and genotype-environment matches for maximizing restoration success. *M. capitata* tends to have lower growth rates in more thermally-tolerant genotypes (Shore-Maggio et al., 2018), but we did not observe the same pattern.

Restoration of coral cover and the resultant biodiversity, structure and function can only occur when corals grow onto the reef and take up once uninhabited space (Omori, 2019). Documenting this process extends beyond simple survivorship and growth and can be monitored through increases in reef structural complexity as a measure of restoration success (Yanovski & Abelson, 2019). The complexity (fractal dimension) of our plots increased immediately upon outplanting, providing context for the detectable increase of rugosity and reef height needed to protect coastal communities and model restoration value (Storlazzi et al., 2019). We also observed a significant interaction between plot type and time, indicating that structural complexity of outplanted plots increased significantly faster through time than control plots. This outcome suggests a positive feedback loop that may be driven by colony morphology and available open space, the latter of which may be a practical tradeoff if outplant density supports attachment. Species-specific traits strongly influence structural complexity and reef fish biodiversity (Darling et al., 2017), so the response of individual corals to changing environmental conditions during outplanting may also prompt morphological changes (Anthony & Hoegh-Guldberg, 2003). Structural complexity of coral reefs often directly relates to reef health, with increases in biomass and coral cover being attributed to higher reef complexity (Graham & Nash, 2013), suggesting that our outplanting impacted overall reef function immediately and through time.

This project represents the first effort in Hawai’i to quantify outcomes of restoration using RRE scores, which demonstrates the effectiveness of our restoration approach. We also found no difference in return based on thermal tolerance, suggesting that even in non-stressful conditions the variation in attachment, survivorship, growth and complexity attributed to this trait has limited practical and financial impacts on restoration. We also show that although the RRE score in Hawai’i was lower than some regions, it was not significantly different from the global average, which is highly variable because environmental conditions, species assemblages and the functional niches they fill are different between regions. While this precludes a ‘threshold’ for evaluating success, establishing baseline RRE scores for restoration practitioners in Hawai’i provides important context at local reefs using Hawaiian corals. These values can be used to compare project-specific goals and outcomes, optimize methods and maximize return in the future.

Coral restoration projects are attractive because they attempt to directly counteract an observed or expected degradation, but a common criticism of restoration is that it will not restore reefs at an ecosystem level and cannot keep up with the rate of reef degradation. Incorporating thermal resilience as an outplant attribute may extend effectiveness, but such conservation efforts should not be viewed as a solution to climate change and serve only as a temporary solution while large-scale mitigation of the root causes are implemented (Boström-Einarsson et al., 2020). Nevertheless, there is wide-spread interest in restoration methods as a means to make short-term improvements to reef biodiversity and coral cover (Bellwood et al., 2019; Boström-Einarsson et al., 2020) especially at small scales and for targeted preservation goals. Our results demonstrate the viability of restoration in Hawai’i, show that baseline tradeoffs to thermal tolerance under non-stressful conditions are subtle, and indicate there is no significant impact on return on effort in proactive restoration. Coral restoration projects have historically been limited in Hawai’i, but are receiving increasing interest from resource managers and community stakeholders who perceive the impending threat from warming oceans and recognize the negative impacts to infrastructure, environment, and social well-being. With proper monitoring programs, research-scale restoration projects like this one can inform locale-specific decisions concerning the implementation of coral restoration and associated methodologies.

## Acknowledgments

We are grateful to the Coral Resilience Lab and Kuleana Coral Reefs for field and laboratory support during this project. Maile Villablanca, Darienne Kealoha and Tahirih Perez assisted with photogrammetry analysis. This work was funded by the National Fish and Wildlife Foundation Award 0381.18.062406 (CD), the Paul G. Allen Family Foundation (CD), and the Schmidt Ocean Institute (HHS). Corals were collected and outplanted with approval from Hawai’i Division of Aquatic Resources permits SAP 2019-48, SAP 2021-06, SAP 2022-14, SAP 2023-06, and SAP 2024-14 granted to National Oceanic and Atmospheric Administration. This is SOEST contribution xxx and HIMB contribution xxx.

## Data Availability

All data and code needed to reproduce this analysis is available at github.com/CarloReef/RWR_Airport.

