## Supplemental Text for "Proactive Coral Reef Restoration Using Thermally Tolerant Corals in Hawai‘i"

**Supplemental Methods**

*Coral Collection, Study Site and Experimental Overview*

In 2018, National Oceanic and Atmospheric Administration (NOAA) divers collected COO for mitigation efforts in response to the grounding of an 83-foot commercial vessel on the fringing reef directly offshore from the Daniel K. Inouye Airport in Honolulu, Oʻahu. Coral colonies were collected from a ~200m x 50m area (~N 21°17.750’, W 157°55.561’) located offshore of the airport. The corals were staged at an *in situ* nursery table located on a sand patch within 100m of the collection site in ~18m of water. In March 2020, we stress-tested biopsies from each colony and identified thermally tolerant and non-tolerant individuals, of which 391 were outplanted. We established 23 plots approximately 6 meters in diameter on a nearby reef slope at a depth ranging from 8.2-14.5 meters over a total area of approximately 725 m^2^ (Figure 1B). A permanent metal pin was placed at the center of each plot and tagged for long-term monitoring using photogrammetry. Plots were randomly designated as control or outplant sites and outplant sites were randomly assigned to higher or lower thermal tolerance corals, detailed below.

*Heat Stress Testing*

COO were tagged and a 5 x ¼” steel rod was glued in a drilled hole in the bottom surface of each colony such that several inches of rod protruded. We sampled 2 biopsies from each colony (~5cm branch or 3cm diameter core) and transported them to the Hawaiʻi Institute of Marine Biology (HIMB). We quarantined biopsy samples at ambient temperatures for a month to allow recovery from sampling stress and to acclimate fragments prior to heat stress exposure. Stress testing was performed in 4 tanks, each ramping from 24 °C to 32 °C over 17 days, held at 32 °C for 7 days, and finally ramped to 33 °C over a day (Figure 1A). This temperature profile approximates an established stress-test which resolves natural bleaching responses in corals (Caruso et al., 2025; Drury et al., 2022). Photos were taken every other day and used to assign a visual bleaching score of each fragment throughout the stress test. Corals were scored on a categorical scale where 3 represented healthy, 2 represented visual paling, 1 represented bleached and 0 represented complete mortality. The bleaching score of each fragment was averaged over the time series (n=13) to obtain a single heat response metric for each sample. To control for tank effects and differences in species-level heat tolerance, samples were grouped by tank and species before being divided into performance quantiles, with approximately half of each group assigned as high or low tolerance (Figure 1) based on performance above or below the median tolerance. After these designations, source colonies were randomly assigned to outplanting plots corresponding to the designation (“high” or “low”) of their respective sample.

*Outplanting*

We randomly designated 17 plots for outplanting (8 “high tolerance” and 9 “low tolerance”) and 6 as “control”. In August 2020, we outplanted colonies in their designated plots, by drilling a ⅜” hole into hard substrate for each colony, removing the identification tag, and injecting two-part epoxy into the hole to glue the steel pin into the reef. Colonies were haphazardly placed within a ~3m radius of the central marker wherever stable, bare substrate was available. Each colony was photographed *in situ* immediately after outplanting and indexed for future identification. No outplanting was performed in control plots.

*Monitoring and 3D Models*

To monitor the reef plots and outplanted coral colonies, we used photogrammetry to produce orthomosaic and digital elevation model (DEM) imagery to assist with mapping and data collection. Baseline imagery was collected in June 2020 before outplanting to enable a comparison of before and after reef complexity data and assist with locating outplants. After outplanting, we performed monitoring quarterly until the second year and bi-annually thereafter. At the time of writing, data collection was complete for 7 timepoints spanning 25 months. Image collection followed the general workflow described in Roach et al. (2021). A spool with a 3.5 m line tethered to a camera was positioned at the center of the plot and used to guide a swimmer in a spiral path to promote full coverage of approximately 28 m^2^ (~3m radius) area of reef. Cameras were set to shoot continuously resulting in an approximate of 600 photos per plot per time point. Photos taken at each reef plot were used to reconstruct orthomosaics and DEMs using Metashape Professional photogrammetry software (Agisoft, LLC). Camera and computer hardware are detailed in Supplemental Table 1.

Each photoset was imported into a single Metashape project file to assist with side-by-side GPS alignment and overlay for data collection. Workflow and reconstruction processing settings were similar to those outlined in Roach et al. (2021)., with the addition of georeferencing the model using AddTools for Metashape’s Calculate GPS Coordinates tool (Settide, LLC). Once the model was georeferenced, each plot’s orthomosaic and DEM imagery were produced and exported for data collection.

*Data Extraction*

To assess initial cover, we randomly selected the pre-outplanting orthomosaics of 10 plots and added 100 random points to each using the Random Points tool in AddTools for Metashape. We then classified the substrate under each point as living coral or not. To collect information about colony level growth, survivorship, and geometric complexity, we identified corals in the orthomosaic imagery and outlined them using the Metashape polygon shape tool. A new Metashape project file for each plot was created separately for data collection to allow feasibility of imagery overlay, batch data collection, and to reduce computer resource usage (i.e. RAM). The orthomosaic and DEM imagery of all time points for each plot were imported into a single Metashape chunk. Using the orthomosaic image, outplants were identified and outlined using the polygon shape tool. Surface area, top-down area, height range, and volume were collected in meters using the DEM Measure tool in AddTools for Metashape. Fractal dimension was calculated at the reef plot scale following (Torres-Pulliza et al., 2020).

To compare the growth and survivorship of coral outplants with corals growing naturally on the reef, the 6 control reef plots were monitored via photogrammetry as described above. To identify and collect data for corals in the control plots, 50 random points within the initial timepoint of each control plot were created on the orthomosaic image using the Random Points tool in AddTools for Metashape. The first 25 colonies located beneath a point at the initial timepoint were selected as control corals. If any part of a coral was underneath or covered by another colony or substrate through any time point, then the coral was not selected as a control.

To gather information about both the photogrammetric reconstruction accuracy and the scale accuracy (linear, area, volumetric), they were calculated using AddTools for Metashape for each plot across all time points (Supplemental Figure 1). All methods described above for collecting measurements of outplants and outplant plots were applied to control corals and control reef plots.

*Statistical Analysis*

All data analysis was performed using R software (2024.04.2+764, (R Core Team, 2018)). To examine the effect of outplanting on plot complexity, we used a paired t-test to compare the fractal dimension of plots before and immediately after outplanting. We used a linear mixed model in the R package *lme4* (Bates et al., 2015) to examine plot complexity over time between control plots and outplant plots [fractal.dimension ~ time * type + (1|plot), where “ type” is control plot or outplant plot]. We used the same modeling approach to examine plot complexity over time between thermal tolerant plots and non-tolerant plots [fractal.dimension ~ time * thermal.tolerance + (1|plot)]. Model residuals were examined and variance assumptions were confirmed; for ease of visualization, we relativized the fractal dimension time series to the initial timepoint, but all analysis was conducted on raw values.

To examine and visualize attachment and survival of outplanted vs control and thermal tolerance levels, we used Cox proportional hazards models implemented using the R packages *ggsurvfit* (Kassambara, A., Kosinski, M., & Biecek, P, 2021), *survival* (Gray, 2002), and *survminer* (Kassambara, A. & Kosinski, M., 2021). First, we evaluated attachment of corals (i.e., retention of corals independent of survivorship) using the survival analysis in a ʻtime-to-eventʻ framework to compare control and outplanted plots [Surv(time,attachment.status) ~ type, where “type” is control plot or outplant plot]. We then right-censored unattached corals and evaluated survivorship of attached corals between control and outplanted plots [Surv(time,survival.status) ~ type, where “type” is control plot or outplant plot]. Next, we evaluated survival of corals between high and low tolerance plots [Surv(time,survival.status) ~ thermal.tolerance]. We used the function pairwise_survdiff to extract significance values between thermal tolerance for each species.

We analyzed coral growth using whole colony surface area metrics determined by photogrammetry as described above. After examining residuals and variance assumptions, we log10 transformed surface area data. We used a linear mixed model to evaluate growth differences between control and outplant corals with individual corals nested within plot included as a random effect to account for repeated measures [surface.area ~ time * type + (1|plot/coral), where “ type” is control or outplant]. We used the same modeling approach to evaluate differences between tolerant and sensitive coral growth rate [surface area ~ time * thermal.tolerance + (1|plot/coral)]. All growth models were visualized using the R package *sjPlot* (Lüdecke D., 2024) from the model outputs.

We used a Relative Return on Effort analysis described in Henry et al. (2023), Howlett et al. (2021), and Suggett et al. (2019) to evaluate the impact of thermal tolerance and provide context for restoration in Hawaii compared to global efforts. Briefly, we calculated arcsine-transformed endpoint survivorship and ln-transformed standardized growth rate of each species for each thermal tolerance grouping and calculated the RRE score. We used a Wilcoxon test to compare RRE scores based on thermal tolerance. We then re-calculated growth and survivorship for each species (without thermal tolerance separation) and compiled our data alongside RRE results from Henry et al. (2023), Howlett et al. (2021), and Suggett et al. (2019). We averaged the RRE score from all other regions and used a one-way t.test to compare the distribution of values from Hawaii to the global mean. We then used a one.way test in R (Welchʻs ANOVA allowing for unequal variances) after exploring model assumptions to evaluate if RRE score varied by region.
