## Supplementary figures and images for "Proactive Coral Reef Restoration Using Thermally Tolerant Corals in Hawai‘i"

### Supplemental Figure

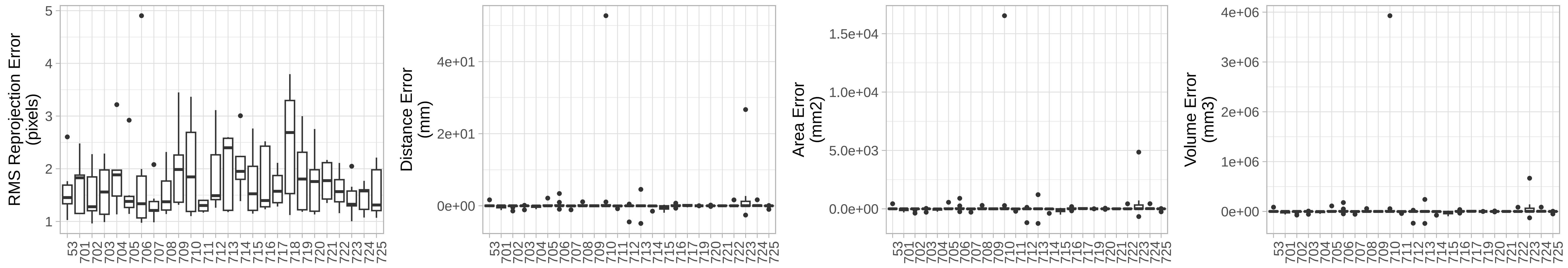
